# Structure of the Native Chemotaxis Core Signalling Unit from E-gene lysed *E. coli* cells

**DOI:** 10.1101/2023.01.30.526190

**Authors:** C. Keith Cassidy, Zhuan Qin, Thomas Frosio, Khoosheh Gosink, Zhengyi Yang, Mark S. P. Sansom, Phillip J. Stansfeld, John S. Parkinson, Peijun Zhang

## Abstract

Motile bacteria detect ambient chemical gradients and control their locomotion via conserved chemotaxis signaling networks, enabling cells to locate nutrients, potential hosts and other important biological niches. The sensory apparatus of the chemotaxis pathway is an array of core-signaling units (CSU) composed of transmembrane chemoreceptors, the histidine kinase CheA and an adaptor protein CheW. Although chemotaxis pathways represent the best understood signaling systems, a detailed mechanistic understanding of signal transduction has been hindered by the lack of a complete structural picture of the CSU and extended array. In this study, we present the structure of the complete CSU from phage E-gene lysed *E. coli* cells, determined using cryo-electron tomography and sub-tomogram averaging to 12 Å resolution. Using AlphaFold2, we further predict the atomic structures of the CSU’s constituent proteins as well as key protein-protein interfaces, enabling the assembly an all-atom CSU model, which we conformationally refine using our cryoET map. Molecular dynamics simulations of the resulting model provide new insight into the periplasmic organization of the complex, including novel interactions between neighboring receptor ligand binding domains. Our results further elucidate previously unresolved interactions between individual CheA domains, including an anti-parallel P1 dimer and non-productive binding mode between P1 and P4, enhancing our understanding of the structural mechanisms underlying CheA signaling and regulation.

## Introduction

Motile bacteria detect ambient chemical gradients and control locomotion via conserved chemotaxis signaling networks, which enable them to seek out nutrients, potential hosts and other important biological niches (1). The chemotaxis network of *Escherichia coli* has served for decades as a model system for the study of sensory signal transduction and motility behaviors (2). In this system, thousands of copies of transmembrane chemoreceptors, the histidine autokinase CheA and adaptor protein CheW form highly ordered structures known as chemosensory arrays (3), which cooperatively integrate the sensory inputs of multiple receptors to regulate the autophosphorylation activity of CheA. CheA, in turn, donates its phosphoryl groups to the diffusible intracellular response regulator CheY to modulate the cell’s flagellar motors.

Core signalling units (CSU) of the chemosensory array contain six receptor dimers, organized as two trimers of dimers (TOD), a single CheA dimer and two CheW adaptor proteins. The CSU is the minimal molecular unit needed to couple CheA autophosphorylation to receptor control (4). The constituent proteins of CSUs have complex, modular structures. *E. coli* has four canonical chemoreceptors (Tsr, Tar, Trg, Tap), known as methyl-accepting chemotaxis proteins (MCPs), that form homodimers of largely helical protomers. MCP molecules contain a periplasmic ligand-binding domain (LBD), transmembrane (TM) four-helix bundle, and cytoplasmic kinase control domain (2). The cytoplasmic domain itself has a membrane-proximal HAMP domain (found in histidine kinases, adenylyl cyclases, MCPs, and some phosphatases) coupled to a long coiled-coil four-helix bundle, the membrane-distal end of which contains the contact surfaces for TOD formation and CSU assembly.

CheA functions as a homodimer in the CSU; each protomer comprises five domains (P1-P5) joined by flexible linkers (5). The P3, P4 and P5 domains form a compact core through the principal dimerization determinant P3. That dimeric core is integrated into the larger CSU via P5 interactions with the receptor tips and CheW. The P4 domain binds ATP and catalyzes autophosphorylation at a histidine residue in the P1 domain. The P1 and P2 domains are attached to the P3-P4-P5 core via long, disordered linkers and mediate phosphoryl group transfer to CheY. The monomeric CheW protein has a fold homologous to the CheA.P5 domain and interacts with P5 to couple CheA to the receptor trimers of the CSU. We note that two additional CheW molecules can associate with the periphery of the CSU and contribute to the formation of hexameric CheW rings in the larger array (6–8). Those CheW molecules are not essential for kinase regulation in the CSU.

High-resolution CSU structures are central to understanding their molecular signaling mechanisms, particularly receptor-mediated kinase control. Toward this end, cryo-electron tomography (cryoET) has played an essential role in revealing that chemoreceptors in diverse microbial species form TODs organized into a hexagonal lattice with identical spacing (9,10). In *E. coli*, this hexagonal packing was shown to extend to the CheA/CheW baseplate region at the receptor tip, where rings containing CheA.P5 and CheW were seen to interlock adjacent CSUs (6,7,11). CryoET studies employing subtomogram averaging (STA) have produced increasingly detailed structures of the CSU in thinner array samples, including those of lysed cells (12), minicells (13), and *in vitro* reconstituted arrays (7,14). Such cryoET density maps have been combined with integrative modelling and molecular simulation techniques to achieve residue-level characterization of CSU structure and dynamics (7,13,14). Nevertheless, shortcomings remain: Reconstructions from lysed cells have so far achieved only modest resolutions (>20 Å) and minicells can still be relatively thick (∼400 nm), which limits the maximum resolution possible. Moreover, although reconstructions of monolayer arrays reached 8 Å resolution, they were formed using soluble receptor molecules that lacked the periplasmic and transmembrane regions.

Here we utilized controlled lysis triggered by induction of a phage lysin to create thin *E. coli* ghost cells containing wild-type chemosensory arrays. We further used STA to obtain a density map of the complete CSU and AlphaFold2 to produce full-length atomic models of the CSU proteins that could be flexibly fitted to the cryoET density map. All-atom molecular dynamics (MD) simulations of the resulting complete CSU model provided new insights into the periplasmic organization of the CSU and on previously unresolved interactions between individual CheA domains, suggesting a new generation of structure-function experiments to elucidate the workings of chemosensory arrays.

## Results

### Imaging native chemosensory arrays in wild-type *E. coli* ghosts

The thickness of native *E. coli* cells (∼1.5 μm) prohibits acquisition of quality cryo-tomograms for high resolution structural analysis of chemosensory arrays. We previously developed a method to produce *E. coli* ‘ghost’ cells just before plunge-freezing (15). Upon induction of lysis gene E from the φX174 bacteriophage, spot lesions form in the cell membranes that release the cellular contents while maintaining the overall integrity of the cell envelop. Vitrification of such samples produces thin, flattened cell ghosts (<200 nm) (Figure 1A). In light microscopy, the *E. coli* ghosts were easily distinguished from unlysed cells due to minimal cytoplasmic GFP-labeled CheY, and had distinct chemosensory arrays as visualized via associated CheY (16) (Figure 1B, arrows). The *E. coli* ghosts were also easy to identify in cryoEM projection images as they appeared semi-transparent. The resulting reconstructed tomograms display well-ordered hexagonal polar arrays about 200-300 nm across with a 12-nm lattice spacing (Figure 1C-D, Supplementary Movie 1).

**Figure 1.**
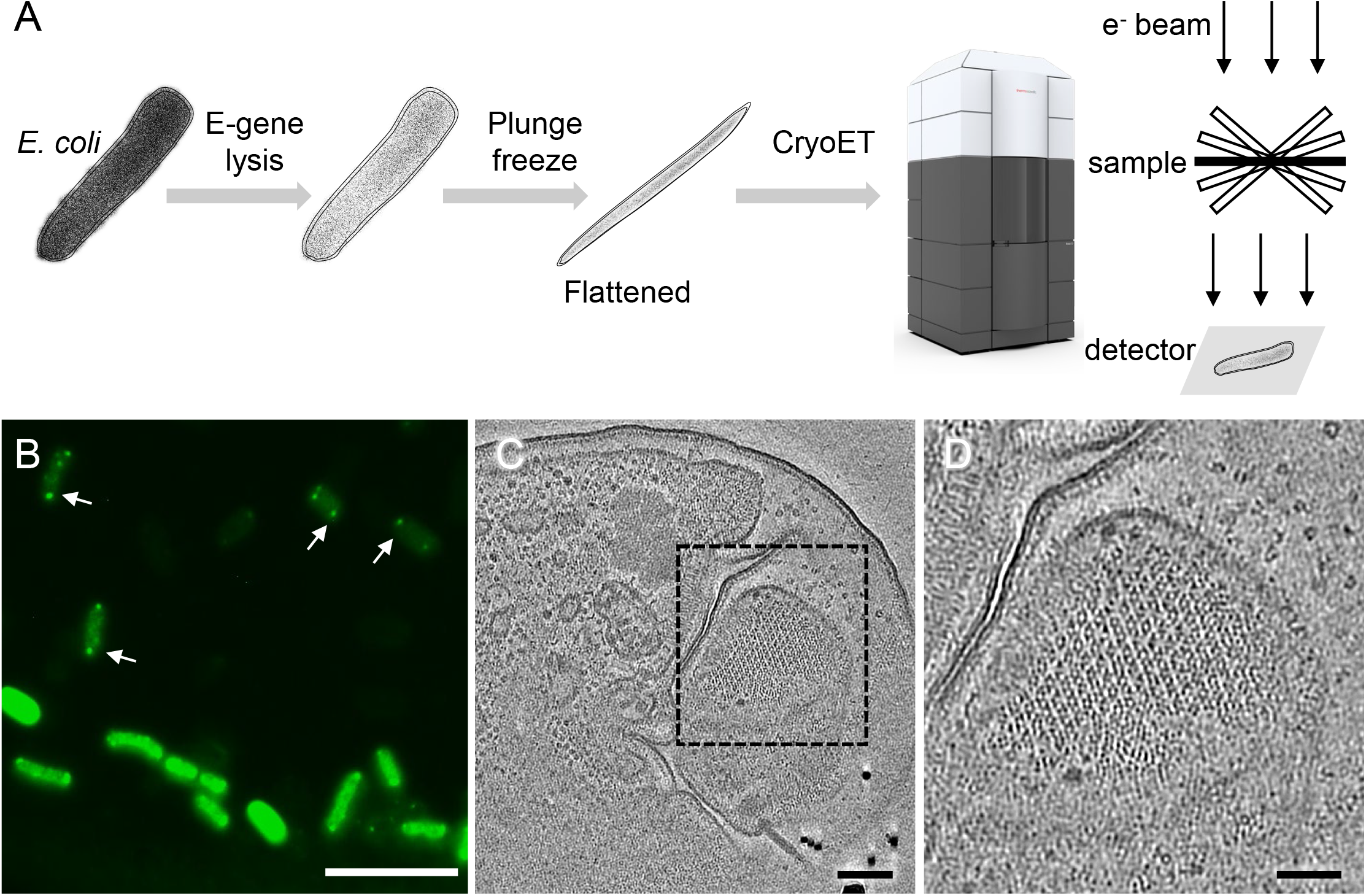
CryoET of native chemosensory arrays in wild-type *E. coli* ghost cells. (A) A schematic diagram of the cryoET workflow used in this study. (B) A fluorescent light microscopy image of GFP-labeled CheY highlights chemosensory arrays (arrows) near the poles of both E-lysed cells (transparent green) and unlysed cells (bright green). (C) A tomographic slice of an E-lysed *E. coli* cell highlighting a patch of the chemosensory array lattice (dashed box). (D) A zoom-in view of the area marked by the dashed box in C. Scale bars, 10 µm in B, 100nm in C, and 50nm in D.

### CryoET STA structure of the complete CSU in *E. coli* ghosts

CryoET data were collected from the E-gene lysed *E. coli* ghosts. The data collection and processing statistics are summarized in Supplementary Table 1, and the cryoET STA process is illustrated in a workflow schematic (Supplementary Figure 1A). A low-resolution structure of the CSU lattice was used for template matching in emClarity (17). The resulting convolution maps (Supplementary Figure 1B-C, Supplementary Movie 2) display cross-correlation peaks for the selection of subtomograms (Supplementary Figure 1D-E). Although, the template-matched subtomograms all contained six receptor trimers in a hexagonal lattice (Supplementary Figure 1A-iii, Supplementary Figure 1E), they were heterogeneous at the CheA/CheW baseplate level and resulted in four structural classes (Supplementary Figure 1A-iv). Subtomograms centered on a CSU trimer (Supplementary Figure 1A-iv, classes with green checks) were selected (Supplementary Figure 1F) and subjected to further refinement (Supplementary Figure 1A-v), from which subtomograms containing a single CSU were iteratively aligned and averaged to give rise to the final CSU map (Supplementary Figure 1A-vi).

The map of the full CSU, determined at an overall resolution of 12 Å (Figure 2A, Supplementary Figure 2A), clearly resolves six full-length chemoreceptor dimers (Figure 2A, red), a central CheA dimer containing distinct regions for P3-P5 (Figure 2A, blue), four CheW monomers (Figure 2A, orange and yellow), and the lipid bilayer headgroups (Fig. 2A, gray). The architecture of CSU at the cytoplasmic side is consistent with the recently reported CSU structure at 16 Å resolution from *E. coli* minicells (13) (Supplementary Figure 3A-C). However, the improved resolution, particularly within the periplasmic region, provides better localization of each protein domain. Both maps show similar splaying of the receptors within each TOD as well as comparable baseplate densities, particularly regarding the shape and orientation of a CheA “keel” density that contains the P4 domain (Supplementary Figure 3C). We also observed additional keel density apparently not attributable to CheA.P4. Comparison of the present CSU structure with the one derived from *in vitro* monolayer arrays at 8 Å (14) shows that the latter structure does not contain corresponding density in this region (Supplementary Figure 3D-G).

**Figure 2.**
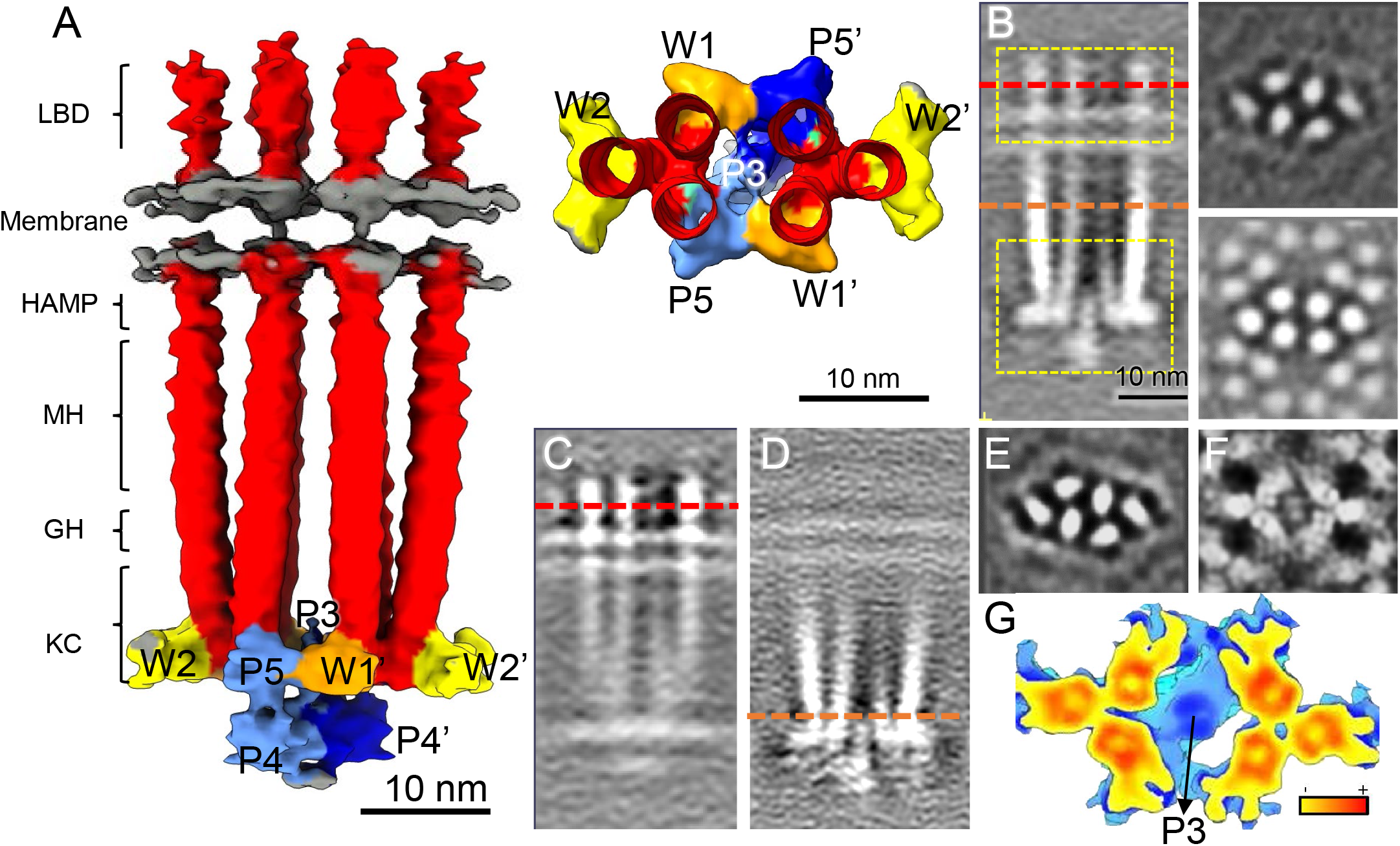
CryoET subtomogram averaging of the core signaling unit (CSU) of native chemosensory arrays. (A) Surface rendering of the CSU density map. Densities corresponding to the receptors are colored in red, CheA dimer in light/dark blue, CheW in yellow/orange, and lipid headgroups in gray. Receptors regions are labelled as follows: ligand binding domain (LBD), HAMP, methylation helix bundle (MH), glycine hinge (GH), and the kinase control region (KC). Individual CheA domains and CheW molecules are labeled accordingly. To the right, a top view of the CSU, from the glycine hinge down, is shown. (B) Sectional side view (left) and top views (right) of the CSU density map. The top views are at the chemoreceptor ligand binding domain (dashed red line) (top right) and at the cytoplasmic domain (dashed orange line) (bottom right), respectively. (C-D) Focused alignment and averaging centered on the receptor LBDs (top dashed yellow box in B) (C) and baseplate region encompassing CheA, CheW and the receptor signaling domain (bottom dashed yellow box in B) (D). (E-F) Sectional views of the averaged maps from C and D at the receptor LDBs (dashed red line in C) (E) and baseplate (dashed orange line in D) (F). (G) An enlarged view of the refined CSU baseplate, colored from blue to red, according to the density value.

Notably the monolayer system utilized truncated CheA molecules not possessing P1 and P2 domains, suggesting that the excess density likely corresponds to these domains. Docking of the molecular model derived from the *in vitro* system (PDB 6S1K) (14) allowed further localization of the excess density, positioning it directly between and below the two P4 domains (Supplementary Figure 3H).

### Focused density refinement of the CSU periplasmic and baseplate regions

Local resolution analysis of the full CSU suggested that the complex was quite flexible in the periplasmic and CheA.P4 regions (Supplementary Figure 2B). We, therefore, carried out a focused refinement of the periplasmic and baseplate regions separately (Figure 2B yellow boxes). Indeed, as the density of each localized region improved during focused refinement, the remainder of the complex became poorly resolved (Figure 2C-D), suggesting considerable structural flexibility between these two regions. The focused refinements also resolved additional structural details in both regions. In the periplasmic-focused map, the ellipsoidal shape of the individual LBDs became prominent and clearly indicated the orientation of each LBD (Figure 2E), a feature not discernible in previous studies. These ellipsoidal densities showed a three-fold symmetry within each TOD, although no such symmetry was applied during refinement.

Similarly, in the baseplate-focused map, the individual α-helices in the receptor signaling domain and CheA P3/P3’ dimerization domains became better resolved (Figure 2F-G). Notably, although the alignment was done just below the glycine hinge region of the receptors (see Figure 2A), the density of the six receptor dimers remained strong from the baseplate to the methylation helix (Figure 2D), indicating that receptors have stable structures throughout the cytoplasmic coiled-coil bundle but increase in flexibility near the HAMP domain. This result is consistent with observations of negatively-stained receptors embedded in nanodiscs, which were often bent just below HAMP (18). In contrast, the glycine hinge region, which evinced signaling-related structural differences in a recent study (12), was relatively stable in the intact wild-type arrays we imaged.

### All-atom model of the complete *E. coli* CSU

Existing high-resolution information on the structures of full-length chemotaxis proteins is sparse. Previous *E. coli* structural studies have largely relied on the use of analogous structures from *T. maritima*, a distantly related organism, either directly or as templates for homology modelling, to assess residue-level details. Recently, AlphaFold2 (AF2) (19) has been shown to produce high-fidelity atomistic structures of proteins and complexes using machine-learning, providing a powerful alternative means of obtaining high resolution structural information, especially in experimentally challenging cases. Accordingly, we used AF2 to construct models of the *E. coli* CSU proteins and sub-complexes, including full-length Tsr and CheA as well as the (CheA.P5/CheW)_3_ and (CheW)_6_ hexameric rings comprising the array baseplate (Figure 3A-C). Overall, the predicted structures are of high quality as assessed by pLDDT, a per-residue confidence score output by AF2, and agree well with existing high-resolution structures and previous modelling results as described below.

**Figure 3.**
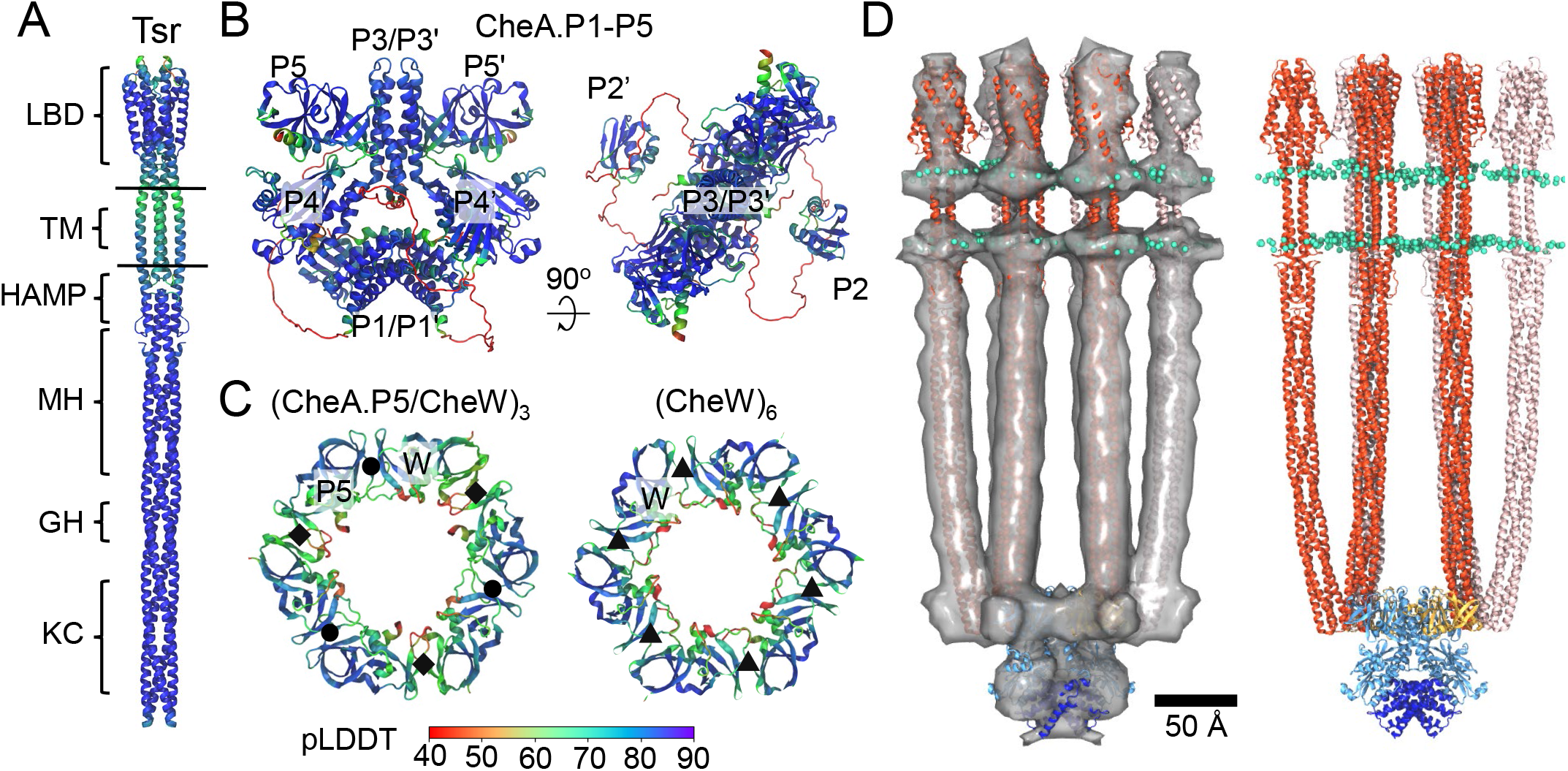
All-atom model of the full-length *E. coli* core signaling unit (CSU). (A-C) AlphaFold2 models of *E. coli* Tsr (A), CheA (B), and intact (CheA.P5/CheW)_3_ and (CheW)_6_ rings (C). Individual domains are labelled as in Figure 2. Models are colored by pLDDT, a residue-based pLDDT confidence score. In panel C, interfaces I, II, and III are denoted by black circles, diamonds, and triangles respectively. (D) MDFF-refined CSU model with (left) and without (right) overlayed density map. Tsr dimers are shown in red/pink, CheA.P3-5 is shown in light blue, CheA.P1 in dark blue, CheW in gold, and lipid headgroups in cyan.

The predicted full-length Tsr structure (Figure 3A) is consistent with X-ray crystal structures of the Tsr cytoplasmic domain (PDB 3ZX6) (20) and APO Tsr periplasmic domain (PDB 2D4U) (21). Additionally, the Tsr transmembrane (TM), control cable, and HAMP regions, which were previously unresolved, closely resemble those seen in a crystal structure of the symmetric APO state of the NarQ sensor kinase (PDB 5JEQ) (22) and agree well with previous modelling results based on TM cross-linking data from *E. coli* Tar (13,23). In particular, the predicted TM bundle displays a central TM1/TM1’ dimer with flanking TM2 helices, as previously proposed for *E. coli* chemoreceptors generally (24), and the critical control-cable segment connecting TM2 to the AS1 helix of HAMP is observed to be helical and kinked as previously suggested based on mutagenesis data (25).

The predicted folds of the individual *E. coli* CheA domains and their inter-domain arrangements (Figure 3B) agree well with existing crystal structures from *T. maritima* (26,27). The P3 and P4 domains adopt a planar arrangement similar to that seen in a crystal structure of a soluble *T. maritima* CheA.P3-P4 construct (PDB 4XIV) (27) and closely resemble the ‘undipped’ CheA conformation deduced from cryoET maps of *in vitro* monolayer arrays (PDB 3JA6 and 6S1K) (7,14). The P5 regulatory domains are rotated slightly compared to the orientation required to form interface I with CheW. Intriguingly, the P1 domains were observed to form a dimer wedged between both P4 domains (Figure 3B), matching precisely to the aforementioned excess density in our cryoET map (Figure 3D). The locations of the P2 domains, on the other hand, vary considerably between AF2 predictions (Supplementary Figure 4). The functional implications of these observations are discussed in more detail below. Finally, in line with the disordered nature of the long, flexible linkers connecting P1 and P2 to one another and P3 (28), AF2 predicts these regions to be entirely unstructured with a low pLDDT score.

To aid in the computational assembly of the CSU, we also utilized AF2 to construct models of key protein-protein interfaces, including the trimeric interface between the Tsr protein-interaction regions, the Tsr/CheA.P5 and Tsr/CheW interfaces, interfaces I and II between CheA.P5 and CheW, and interface III between two CheW molecules. In the case of interfaces I and II, the AF2-predicted structure of the (CheA.P5/CheW)_3_ ring (Figure 3C) agrees well with the analogous *T. maritima* crystal structure (PDB 4JPB) (29), as well as *in vivo* cross-linking studies in *E. coli* (30). Additionally, interface III is in good agreement with molecular models derived via analogy with the (CheA.P5/CheW)_3_ ring and validated with *in vivo* cross-linking data (8,31). The constituent models were assembled using the predicted interfaces and our cryoET map to produce a preliminary model of the CSU, which was subsequently refined using molecular dynamics flexible fitting (MDFF) (32) (Supplementary Figure 5). Finally, the refined CSU model was embedded in an atomistic lipid bilayer (Figure 3D) with a composition chosen to mimic that of the *E. coli* inner membrane (33). Full details regarding the modelling procedures are provided in the Methods.

### Neighboring receptors form transient interactions within the periplasmic space

The improved characterization of MCP LBD orientations in our cryoET STA map permitted positioning these domains within the full CSU structure with added confidence (Figure 4A). To test the robustness of the obtained fits, we conducted a series of MDFF simulations in which the receptor LBDs were rotated by 30°, 60°, or 90° from their initial positions. Indeed, in each simulation, the LBDs returned to their original fitted orientations. We additionally noted potential interactions between neighboring LBDs as they rotated past one another. We therefore decided to carry out an all-atom simulation of the CSU to explore the possibility of specific inter-receptor interactions within the periplasmic space. The resulting trajectory (Supplementary Movie 3) identified interactions between two, co-planar clusters of charged residues on the Tsr LBDs, involving K99, E102, and K103 on helix 2 (cluster 1) as well as E124, K126, R127, and D130 on helix 3 (cluster 2) (Figure 4B). In addition to interactions between receptors within the same TOD, we also observed interactions between the two TODs (Figure 4C). Given the symmetry of the extended array architecture, especially in the periplasmic space (31), such interactions presumably also occur between receptors from neighboring CSUs. Notably, however, the observed interactions were transient (<10 ns) and did not form a single, long-range pattern across the CSU on the timescale of our simulation. Rather, these interactions appear to form a pseudo-lattice by which the Tsr LBDs roughly maintain their relative orientations and inter-domain distances.

**Figure 4.**
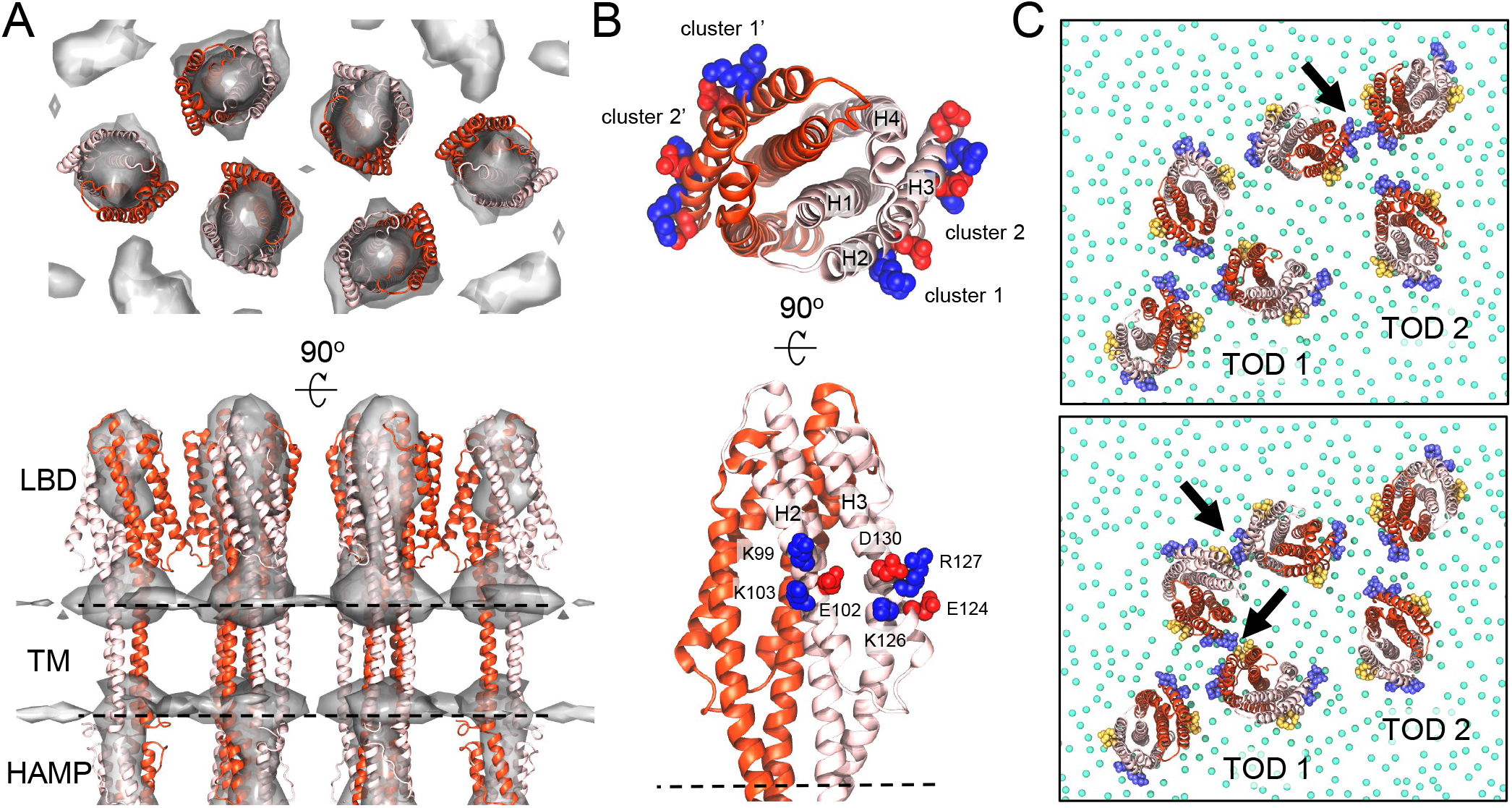
Periplasmic inter-receptor interactions within CSU. (A) Top and side views of the periplasmic region of the refined CSU model, involving the receptor ligand binding domain (LBD), transmembrane (TM) bundle, and HAMP domain, overlayed with the cryoET STA map (shown in Figure 2C&E). Receptor monomers are differentiated using light and dark red. The location of the membrane, not shown for clarity, is illustrated with dashed lines. (B) Top and side views of a single Tsr LBD, highlighting the two clusters of residues observed in an MD simulation to form inter-receptor interactions. Individual residues within each cluster are labelled and shown as VDW spheres in red (acidic) or blue (basic). LBD helices are labelled H1-4. (C) Snapshots from an MD trajectory illustrating interactions (arrows) between receptors within the same trimer-of-dimers (TOD) (top) and between TODs (bottom). For clarity, residues forming clusters 1 and 2 have been coloured yellow and purple respectively and only the headgroup of each lipid is shown (green spheres).

### CheA.P1 forms an anti-parallel dimer that interacts non-productively with CheA.P4

The central P1 dimer predicted as part of the full-length CheA structure by AF2 describes well the excess density between the P4 domains observed in our map (Figure 5A), which was previously noted within a cryoET map from *E. coli* minicells, but could not be assigned atomic structure (13). Moreover, the size and shape of the P1 dimer suggest it alone accounts for this excess, in line with the variable placement of the P2 domains by AF2. Given experimental findings demonstrating that the presence of P2 is not strictly required for effective kinase regulation and chemotaxis (34), these observations suggest that P2 does not form specific interactions within the rest of the CheA core. The P1 domains themselves are predicted to interact in an anti-parallel fashion mediated by P1 helices A and B, such that the substrate histidines face away from the P4 catalytic site and toward the P1/P1’ interface (Figure 5B). The interaction between P1 and P4 is primarily mediated by a string of oppositely-charged residues residing on helix D and the C-terminus of helix A in P1 and the ⍺3 helix in P4 (Figure 5C). Additionally, there exist interactions between the N-terminus of P1 helix A and the N-terminus of the P3 dimerization bundle, particularly between the previously noted P1 residues D14 and E18 (35) and P3 residue R265, which has been shown to play a vital role in CheA function (7,36).

**Figure 5.**
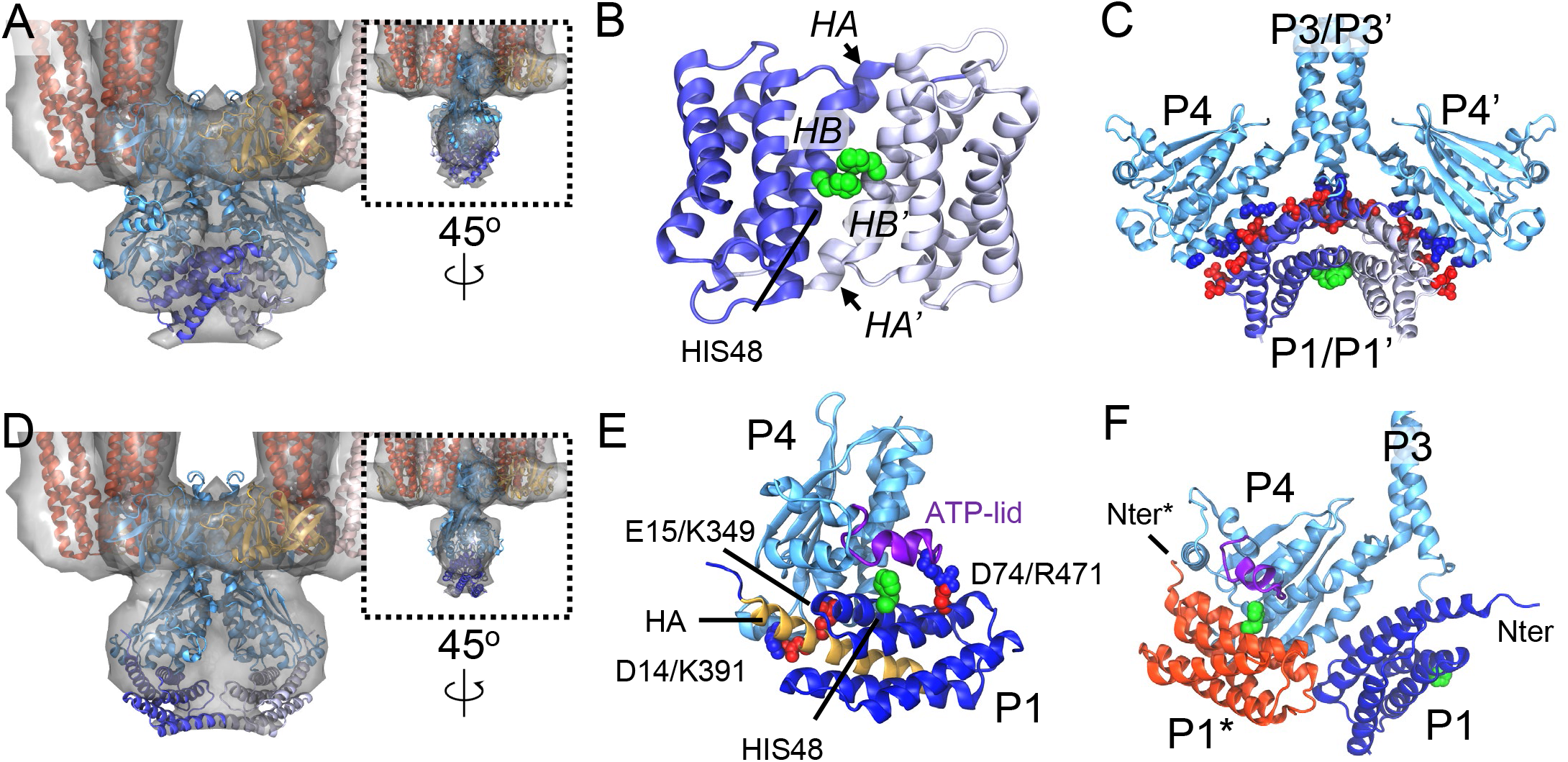
CheA.P1/P1 and CheA.P1/P4 interactions within the CSU. (A) Baseplate region of the MDFF-refined CSU model with density map overlayed, illustrating the fit of the AF2-predicted P1 dimer bound non-productively to P4/P4’. (B) Anti-parallel P1 dimer structure predicted by AF2 for full-length CheA shown in isolation. The Individual P1 monomers are depicted in light and dark blue color and the substrate histidines are colored green and shown in VDW representation. The A and B helices mediating the dimer interaction are labeled for each monomer. (C) Predicted interaction interface between P1 and P3-P4. Acidic and basic residues forming potential inter-domain salt bridges are shown in red and blue respectively. Note, the surrounding regions of the CSU model has been removed for clarity. (D) Baseplate region of the CSU in which a model of the dipped CheA conformation bound productively to P1 has been docked into the overlayed density map. (E) Predicted interaction between CheA.P1 and CheA.P4 for the productive transfer of phosphate from bound ATP (not shown) to HIS48 (green VDW spheres). Residues stabilizing the interaction are labelled and shown as VDW spheres in red (acidic) or blue (basic). CheA.P1 and CheA.P4 are shown in dark and light blue respectively. P1 helix A and the ATP-lid are shown in gold and purple respectively. (F) Overlay of nonproductive (blue) and productive (red) P1/P4 binding modes. The N-terminus of each is labelled with an asterisk to denote the productive binding mode.

In line with this predicted P1/P4 binding surface, a previous NMR analysis of *T. maritima* CheA, designed to monitor the binding of free P1 to a soluble P3/P4 construct, showed large chemical shifts on the P1 D helix as well as on the P4 ⍺4 helix, a region distal to the catalytic site, leading to the proposal of a non-productive P1/P4 binding mode (37). In addition, sizable changes in chemical shifts were also reported on the P1 A and B helices, which form interactions with both P4 and the neighboring P1 monomer in our predicted structure, while the P1 C and E helices, which showed little change in chemical shifts, do not form any inter-domain contacts. In contrast, the predicted CheA structure primarily shows the P4 ⍺3 helix mediating non-productive P1 interactions, whereas the NMR analysis suggested the P4 ⍺4 helix fulfils this role. It’s possible, however, that this discrepancy may arise due to slight alterations in the relative orientations of the P3 and P4 domains due to the soluble and truncated nature of the CheA constructs used in the NMR analysis.

Recently, Muok and co-workers also observed close interactions between the P1 domains within *T. maritima* ‘foldon’ complexes, especially in the inhibited state, and further reported a crystal structure of a *T. maritima* P1 dimer (38). Like the AF2-predicted *E. coli* P1 dimer, this structure showed the substrate histidines facing toward the dimer interface in a non-productive fashion, however, the P1 domains showed a parallel arrangement as opposed to an anti-parallel. Given that the parallel and anti-parallel P1 dimers appear to match the PDS data reported by Muok *et al*. equally well, we asked whether this change might be attributed to differences in P1 sequence or the result of severing P1 from CheA.P3-P5. We therefore used AF2 to (1) predict the structure of two *E. coli* P1 domains alone and (2) predict the structure of the full-length *T. maritima* CheA dimer. In the first case, the resulting prediction showed a parallel *E. coli* P1 dimer very similar to the reported *T. maritima* crystal structure (Supplementary Figure 6), while in the second case, the full-length *T. maritima* CheA structure showed an anti-parallel dimer as in the full-length *E. coli* structure (Figure 5B). We therefore suggest that full-length CheA leads to important interactions between P1 and P4 that favor an anti-parallel P1 interaction in native complexes.

### Dipped CheA.P4 accommodates a single productively bound P1

Previously, we predicted the existence of a “dipped” P4 conformation using molecular simulations of the *T. maritima* CSU (7), which we also subsequently identified in a sub-nanometer cryoET map of the *E. coli* CSU from *in vitro* monolayer arrays (14). Recently, multiple experimental studies have lent strong evidence to the existence of such a conformation and demonstrated its importance for the CheA catalytic cycle (5,36,39). Although, the present resolution prevents explicitly isolating distinct P4 conformations within our cryoET map, a rigid docking of our dipped CheA model produces a configuration that is also consistent with the density (Figure 5D). However, the dipped P4 domains, which are rotated downward and toward one another, considerably reduce the volume attributable to the P1 dimer and prevent the formation of predicted stabilizing interactions with P4, suggesting that the postulated non-productive binding of P1 is not compatible with a dipped P4 conformation. We therefore explored what implications this conformational change might have on the productive binding of P1 to P4.

Previous molecular docking (40) and mutagenesis (35) studies have led to models for the productive interaction of P1 with P4, which involve primarily P1 helices A and B oriented in such a way as to permit the highly-constrained spatial requirements for chemical transfer of the gamma-phosphoryl group from bound ATP to the substrate histidine on P1 helix B (41). Additionally, a previous crystal structure of *T. maritima* CheA.P4 bound to an ATP analog showed that the so-called ATP lid, a typically unstructured span of ∼20 residues near the nucleotide binding site, adopted an ordered conformation, thereby altering the probable interface for productive P1 binding (42). This observation led to the proposal that folding of the ATP lid is a prerequisite for productive P1 binding, a notion which has been further supported by recent kinetic analyses of CheA activation suggesting an ordered sequential mechanism (43).

To assess the implications of these observations in the context of our CSU map and model, we constructed a homology model of *E. coli* P4 with a folded ATP lid based on PDB 1I58 and used HADDOCK (44) to carry out an extensive exploration of potential P1/P4 interactions, suitably restraining the distance between the substrate histidine and gamma-phosphoryl group to encourage productive binding orientations. The top-ranked prediction (Figure 5E) agrees very well with the previously proposed *T. maritima* model (40) and shows interactions mediated primarily by P1 helices A and B, including salt bridges between residues D14/K391 and E15/K349, as well as a potential salt bridge between D74 on P1 helix C and R471 on the P4 ATP lid. Thus, P1 helix A appears to play an important role in both productive and non-productive P1 binding (Figure 5F), although the N-terminus faces inwards towards the P3 bundle in the nonproductive binding mode and is outward facing in the productive binding mode. Notably, the proximity of the productive and non-productive P1 binding modes (Figure 5F), further suggests there may be some degree of steric hindrance to simultaneous binding.

We next used the predicted productive P1/P4 binding mode to map P1 onto the dipped CheA structure (Figure 5D). While the resulting P1 domains mostly fit within the envelope of residual CheA density, compared to the non-productive P1 dimer they leave a considerable volume of density directly below the P3 bundle unfilled. Moreover, unlike the non-productive binding of P1 to undipped P4, which poises the P1 domains to bury a sizable surface area through dimerization, the productively bound P1 domains do not. Rather they face one another through a small cross-sectional area formed by the turns connecting helices A with B and C with D. Given the known conformational flexibility of the P4 domains, especially upon P4 dipping (14), the observed orientations would likely lead to severe steric clashes between two productively-bound P1 domains. We suggest, therefore, that it is unlikely that specific P1/P1 interactions or dimerization play a role in its productive interaction with P4.

## Discussion

In this study, we’ve reported the structure of the chemotaxis CSU from E-gene lysed *E. coli* cells at an overall resolution of 12 Å by cryoET STA. Using AlphaFold2, we’ve further predicted the full-length structures of the CSU’s constituent proteins as well as key protein-protein interfaces, enabling the assembly an atomistic CSU model, which we conformationally refine using our cryoET data. The improved resolution in the periplasmic region allowed for a more precise localization of the positions and relative orientations of the receptor LBDs. An MD simulation of the CSU model in an atomistic lipid bilayer permitted us to directly observe the local diffusion of receptors within the membrane for the first time and identified clusters of charged residues forming interactions between neighboring receptor LBDs. In line with these observations, previous *in vivo* work using a Tar-only *E. coli* strain observed that the efficiency of disulfide cross-linking between residues in periplasmic helix 2 and 3 varied as a function of methylation state and exposure to the chemoattractant aspartate (45). However, analysis of the cross-linking results was based on a crude pictures of CSU structure and array architecture, potentially limiting functional interpretations. Our present model and observations therefore provide a high-resolution structural basis for designing experiments to test for concerted, signal-dependent changes in the periplasmic space.

Additionally, integration of AlphaFold2 predictions of full-length CheA into our cryoET density map, allowed us to incorporate the P1 domains into the CSU model as an anti-parallel dimer, non-productively bound between the P4 domains. As previously noted, this prediction is in line with findings in *T. maritima*, which suggest both the existence of a non-productive P1-P4 binding mode and P1 dimerization (37,38). Moreover, previous *in vivo* cryoET images of *E. coli* CSUs in distinct signaling states imposed by mutagenesis showed that the overall volume of CheA density tended to increase in the kinase-OFF state, suggesting that regulation of P1 and P2 mobility might be a key feature of kinase control (46). Along these lines, the non-productive P1/P4 contact interfaces presented here require both P4 domains to adopt an undipped conformation, which we previously showed to be stabilized by direct P4-P5 contacts (14). The P1 dimer may therefore act as a wedge to further reduce P4 mobility and CheA dynamics overall in the kinase-OFF state. However, our results suggest that P2 is not integral to this ordered state and thus its mobility is unlikely to be directly regulated.

Studies involving the kinetic analysis of CheA autophosphorylation using liberated P1 domains, on the other hand, have discounted the role of P1 sequestration in CheA regulation, proposing that receptors primarily mediate kinase regulation by altering the apparent rate constant of autophosphorylation through control of ATP binding (43,47,48). Nevertheless, our present observations can be reconciled with such a picture if, for instance, ATP binds preferentially to either the undipped or dipped P4 conformation, thereby allowing receptors to indirectly mediate ATP binding through the regulation of P4 dynamics, which could be in turn affected by P1 dimerization and sequestration. It has been previously noted that the undipped P4 conformation situates the unfolded ATP-lid near the P5 regulatory domain where it can form stabilizing interactions (5,14). One possibility therefore is that such interactions discourage the folding ATP-lid, preventing stabilization of ATP binding and the formation of a suitable interface for productive P1 binding (40,42). Another possibility is that differences in the P3-P4 and P4-P5 linker conformations, which have previously been shown to be critical for kinase function and coupled to P4 dipping (39,49–51), might induce subtle changes in the ATP-binding pocket that increase ATP binding affinity for the dipped P4 state. Given that P4 dipping would also destroy the interfaces required to bind the non-productive P1 dimer, such a conformational change should ultimately lead to disassociation of the P1 dimer, freeing it for productive binding to P4.

Moving forward, the use of strategic functional mutations combined with the improved resolutions enabled by cryoET analysis of E-lysis and minicell constructs will provide a promising tool to reveal critical details of signaling mechanisms such as kinase regulation. The overall similarity between present CSU map and that obtained from *E. coli* minicells (13), suggests that insights from both contexts may be transferrable. Additionally, our atomistic CSU model will greatly enhance the potential for the investigation of signaling mechanisms using molecular simulation, providing a unique way to obtain high-resolution mechanistic insight.

## Materials and Methods

### Bacterial strains, plasmids, cell culture and cryoEM sample preparation

The pRY100 plasmid, carrying the phage φX174 lysis E gene under a tacP promoter and a lacIQ repressor control (52), was transformed into the wild-type *E. coli* K-12 strain RP437 cells using a standard protocol (53). For fluorescence imaging, cells were also transduced with a CheY-GFP construct. Cells were grown in the TB broth (1% Tryptone and 0.4 NaCl, pH 7.0) supplemented with 100 µg/ml ampicillin. E-gene expression was induced by addition of 0.5mM isopropyl β-D-thiogalactopyranoside (IPTG) at OD_600_ of 0.6 as previously described (15). 10 min after induction, an aliquot of 4 µl culture, mixed with 1 µl fiducial gold beads (10 nm size), was applied to glow-discharged Quantifoil R2/2 grids and vitrified with a GP2 plunge-freezing device (Leica).

### Cryo-ET data acquisition

The cryoET titled series data were collected using ThermoFisher Tatian Krios G3 instrument, with a K2 summit detector at a pixel size of 2.8 Å and a K3 summit detector at a pixel size of 2.1 Å (see Supplementary Table 1). Data were collected using the SerielEM software (54,55).

### Cryo-ET reconstruction and subtomogram averaging

Motion correction was performed using MotionCor2 (56). The data collected by the K3 detector were rescaled to 2.8 Å/pixel by Fourier cropping in MotionCor2. The tilt series were aligned using fiducial gold beads in Etomo (57). The aligned tilt series were then imported into emClarity (17) for CTF correction and WBP reconstruction. About 15% of the analyzed tomograms contained arrays of CSUs. A total of 33 array-containing tomograms were selected for cryoET subtomogram averaging.

Template matching as implemented in emClarity was used to for subtomogram particle picking. The position and orientation of picked particles were plotted for visual inspection and bad particles were manually removed. The resulting subtomograms were then aligned and classified according to the workflow described in Supplementary Figure 1. A total of 950 CSU trimers from the K2 dataset processed using emClarity initially yielded an average structure at 8.8 Å resolution but with a strong preferred orientation. A second dataset was collected with a K3 detector and additional subtomograms containing side views were added. In addition, single CSU subvolumes were extracted from the CSU trimers through symmetry expansion and subjected to further alignment and refinement using i3 (58).

### Molecular modelling and simulations

Atomic models of the CSU’s constituent proteins and interfaces were predicted using AlphaFold2 (19) via the ColabFold (59) *AlphaFold2_advanced* notebook at https://github.com/sokrypton/ColabFold. The CSU model was assembled via rigid docking of the component models using UCSF ChimeraX (60) and conformationally refined through a series of symmetry-restrained Molecular Dynamics Flexible Fitting (MDFF) simulations (32). All MDFF simulations were carried out in NAMDv2.14 (61,62) and performed in the NVT ensemble at 300 K with a coupling factor of 0.3 applied to the protein backbone. Additional harmonic restraints were applied during fitting to prevent the loss of secondary structure as well as the formation of cis-peptide bonds and chirality errors. The stereochemistry of the entire CSU model was then refined using ISOLDE (63) in ChimeraX and validated using MolProbity (64).

To prepare the refined CSU model for molecular dynamics (MD) simulation, an atomistic lipid bilayer consisting of 70% PVPE, 20% PVPG, and 10% cardiolipin was assembled around the transmembrane region using CHARMM-GUI (65). The resulting structure was then solvated with TIP3P (66) water molecules and 150 mM NaCl and subjected to conjugant gradient energy minimization followed by a series of equilibration simulations conducted as follows. First, solvent and lipids were permitted to relax while the full protein was harmonically restrained following the multi-step equilibration protocol recommended by CHARMM-GUI. Next, restraints on the protein structure were slowly removed over a span of 10 ns by lowering the associated spring constants in increments of 2 ns. Finally, a 120-ns production simulation without restraints was carried out with analyses being conducted on the last 100 ns. All MD simulations were performed using NAMD v2.14 (61,62) and the CHARMM36 force field (67). Production simulations were conducted in the NPT ensemble with conditions maintained at 1 atm and 310 K using the Nosé–Hoover Langevin piston and Langevin thermostat respectively. The r-RESPA integrator scheme was employed with an integration time step of 2 fs and SHAKE constraints applied to all hydrogen atoms. Short-range, nonbonded interactions were calculated every 2 fs with a cut-off of 12 Å; long-range electrostatics were evaluated every 6 fs using the particle-mesh-Ewald method.

## Data availability

All data needed to evaluate the conclusions in the paper are present in the paper and/or the Supplementary Materials. The cryoET STA map of the full CSU has been deposited in the EMDB under accession code EMD-15641. The cryoET STA maps of the CSU periplasmic and baseplate regions after focused refinement and classification have been deposited in the EMDB under accession code EMD-15643 and EMD-15642, respectively. Atomic coordinates for the associated CSU model are deposited in the PDB under accession code 8C5V.

## Acknowledgements

We are grateful to Drs Tao Ni, Jiwei Liu, Luiza Mendonca, Loic Carrique and James Gilchrist for support; We acknowledge Diamond for access and support of the CryoEM facilities at the UK national Electron Bio-Imaging Centre (eBIC, proposal NT21004, CM22941 and BI22910), funded by the Wellcome Trust, MRC and BBSRC. Computational aspects of this research were supported by the Wellcome Trust Core Award Grant Number 203141/Z/16/Z and the NIHR Oxford BRC. This project also made use of computational resources on HPC granted via the UK High-End Computing Consortium for Biomolecular Simulation, HECBioSim (http://hecbiosim.ac.uk), supported by EPSRC (grant no. EP/R029407/1). This work was further supported by the UK Wellcome Trust Investigator Award 206422/Z/17/Z, the UK Biotechnology and Biological Sciences Research Council grant BB/S003339/1, and the ERC AdG grant (101021133).

## Author contributions

P.Z. conceived the research and designed the experiments. Z.Y. prepared samples. Z.Q., T.F. and Z.Y. collected cryoET data. Z.Q. and T.F. performed tomography reconstruction, subtomogram averaging and analysis. C.K.C. performed and analyzed the modelling and MD simulations, supervised by M.S.P., P.J.S. and P.Z.. C.K.C, Z.Q. and P.Z. analyzed the data. C.K.C., Z.Q., J.S.P. and P.Z. wrote the paper with support from all authors.

## Supplementary materials for

**Supplementary Table 1.** CryoET data collection and structure determination of CSU from native *E. coli* cells.

| Data acquisition |  |  |  |
| --- | --- | --- | --- |
| Dataset | Titan Krio (K2) |  | Titan Krio (K3) |
| Voltage (keV) | 300 |  | 300 |
| Detector | Gatan Quantum K2<br>Direct Electron Detector |  | Gatan Quantum K3<br>Direct Electron Detector |
| Energy-filter | Yes |  | yes |
| Slit width (eV) | 20 |  | 20 |
| Super-resolution mode | no |  | yes |
| Pixel size (Å/pixel) | 2.8 |  | 2.1 |
| Defocus range (µm) | -2 to -3 |  | -0.5 to -3 |
| Acquisition scheme | Dose-Symmetric, -60/60,<br>3° step, group 3 |  | Dose-Symmetric, -60/60,<br>3° step, group 3 |
| Total Dose (electrons/Å²) | 120 |  | 120 |
| Number of Frames | 8 |  | 10 |
| Number of Tomograms | 21 |  | 12 |
| Structure determination |  |  |  |
| No. of subtomograms | 3596 |  | 1509 |
| Resolution (Å) | Full CSU | periplasmic | baseplate |
|  | 12.0 | 12.5 | 11.0 |
| Data deposited | EMD-15641<br>PDB-8C5V | EMD-15643 | EMD-15642 |

**Supplementary Figure 1.**
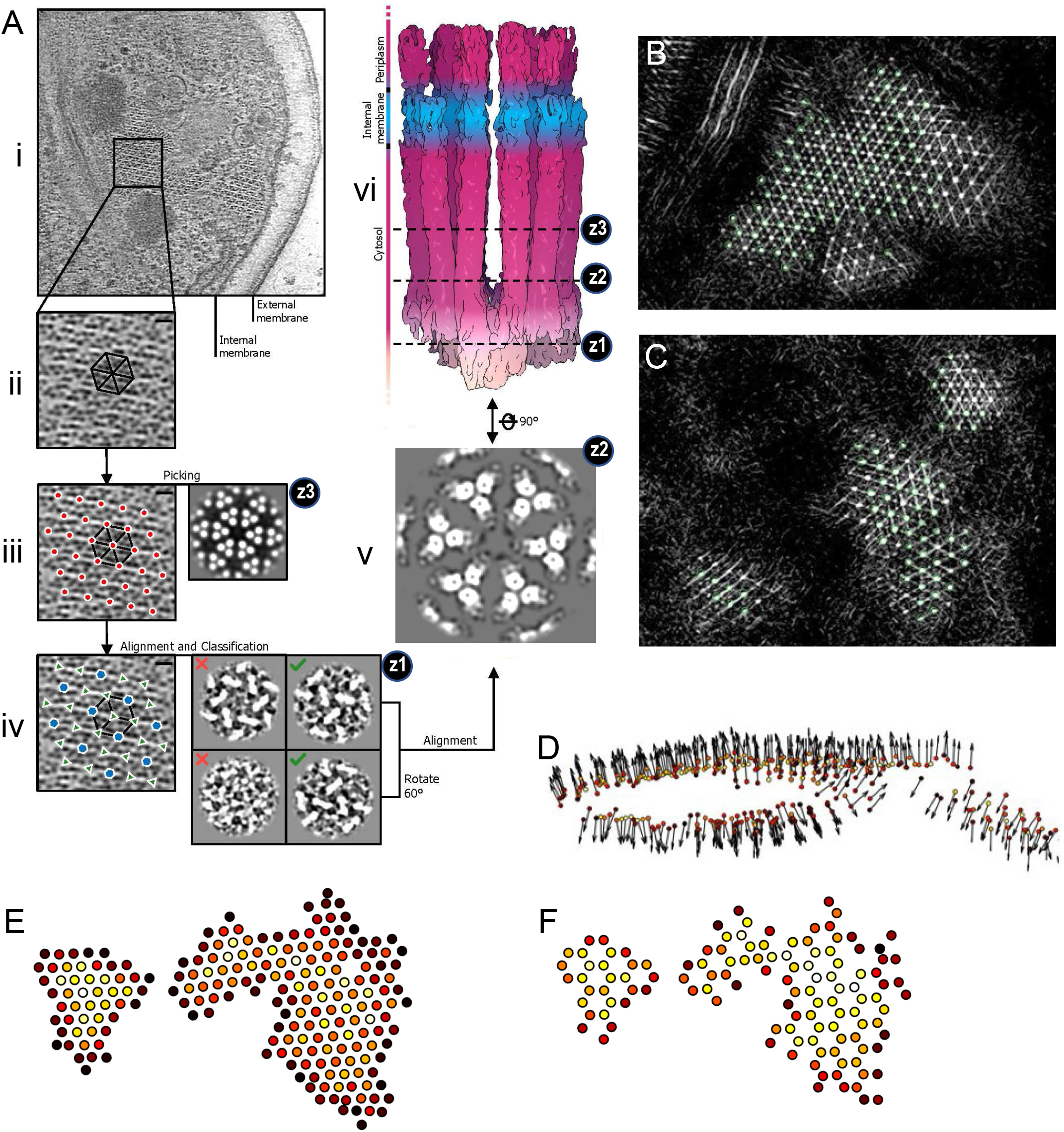
CryoET STA workflow for the CSU trimer. (A) Overview of the cryoET STA workflow. Z1 and Z2 indicate the positions of cross-sections. Steps are indicated as i) raw tomogram, ii) template matching, iii) subvolume picking (the average is shown on the right), iv) alignment and classification (four classes are shown on the right), v) average of CSU trimer, and vi) symmetry expansion and final average of a single CSU. (B-C) Two representative sections of the same conversion map overlapped with picked subvolumes (green circles). (D) Orientations of the picked subvolumes plotted with arrows. (E) Positions of subvolumes after picking in A (iii), where receptor signal dictates. (F) Positions of subvolumes after classification in A (iv), where CSU trimer classes are selected (green checks). The colors indicate cross correlation values, from lowest (black) to highest (white).

**Supplementary Figure 2.**
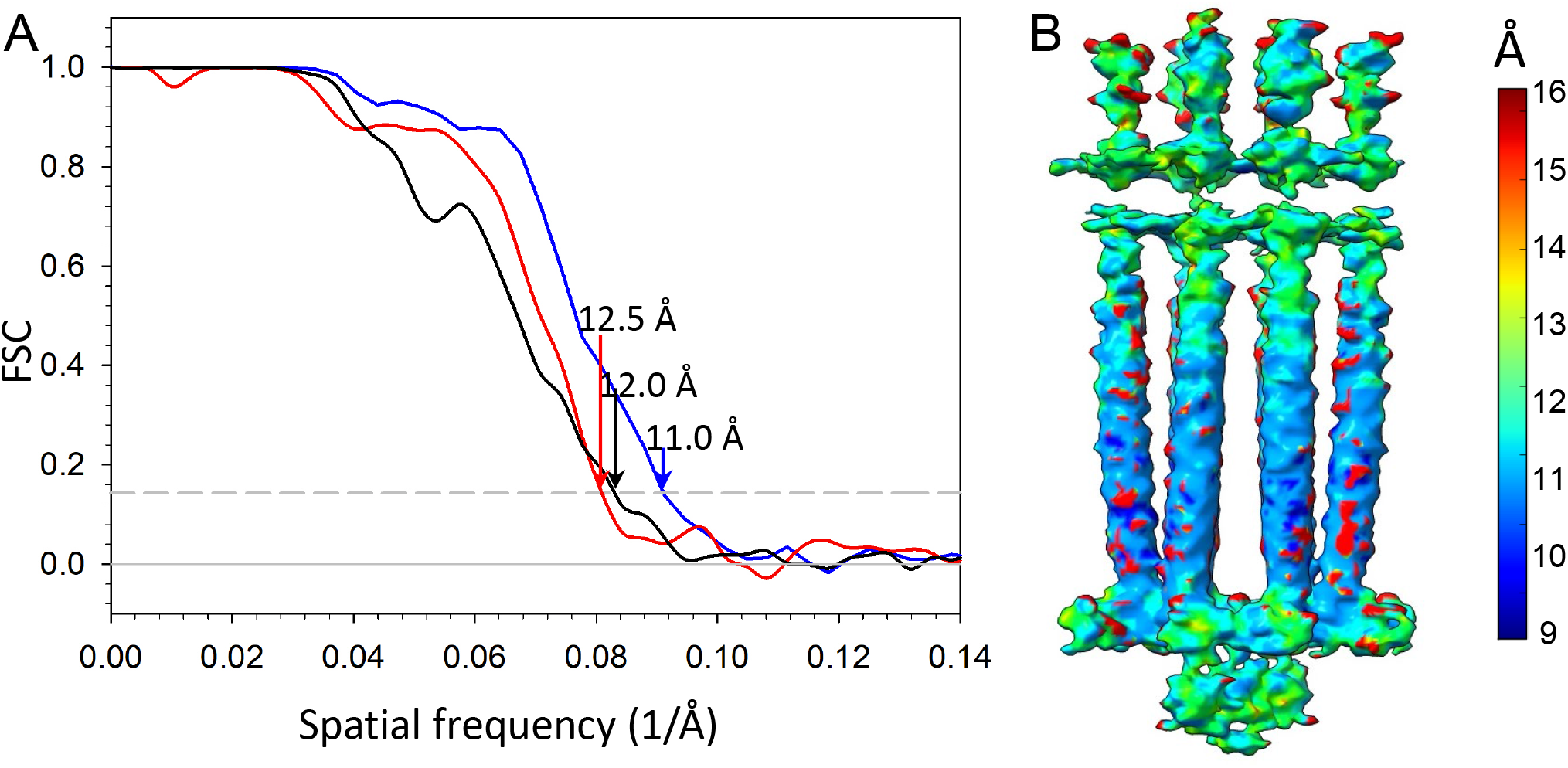
Fourier Shell Correlation (FSC) of the CSU subtomogram average. (A) FSC plots of the overall CSU (black), chemoreceptor ligand binding domains (red), and baseplate region (blue). (B) Local resolution map of the overall CSU, colored from blue to red as indicated in the key.

**Supplementary Figure 3.**
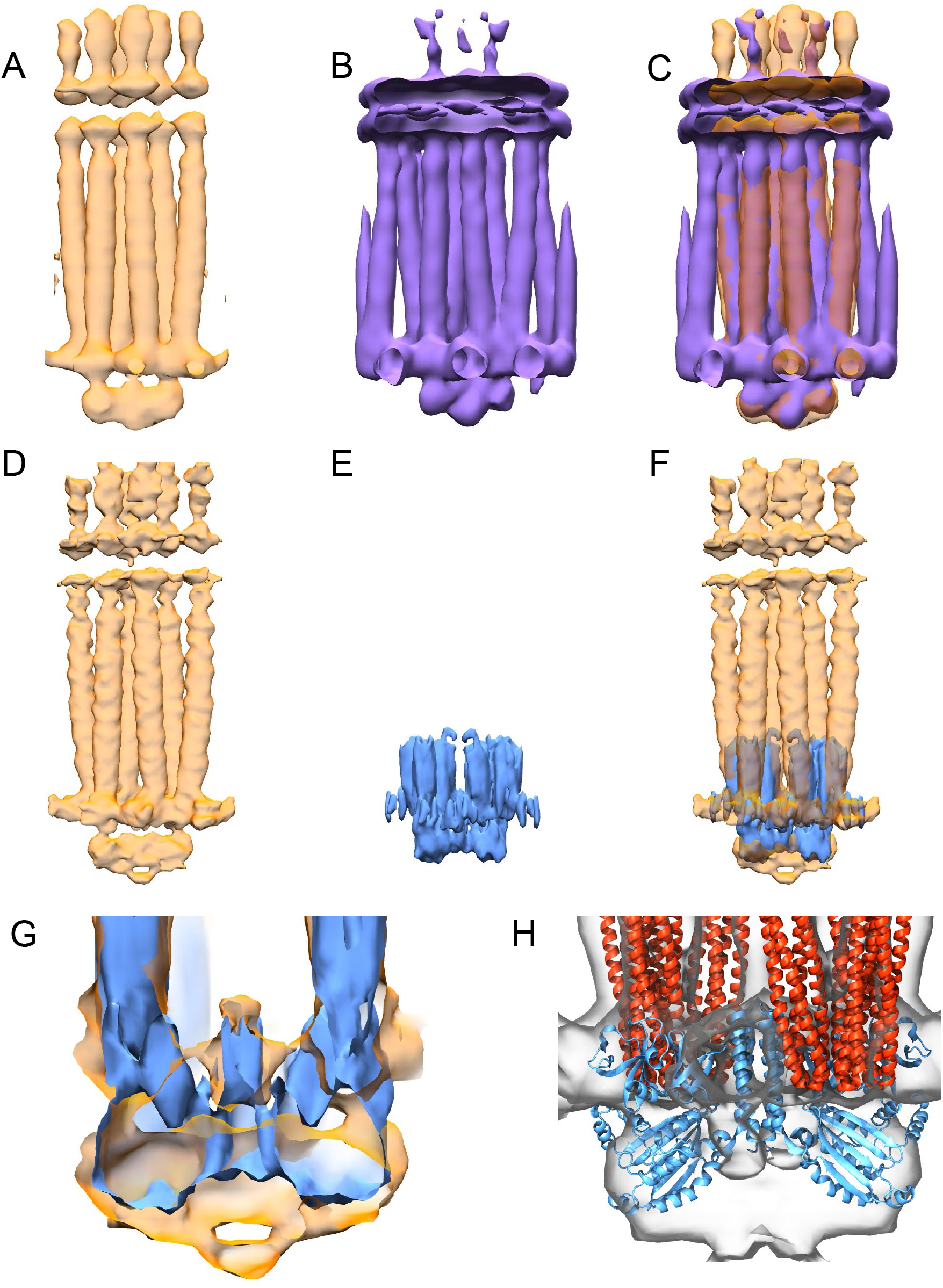
Comparison of CSU maps. (A) CSU map from *E. coli* ghost cells, low-pass filtered to 16 Å resolution. (B) CSU map at 16 Å resolution from *E. coli* minicells (EMD-10160). (C) Overlay of A and B. (D) CSU map from *E. coli* ghost cells, median filtered at 12 Å resolution. (E) CSU map from *E. coli* 4Q monolayer arrays (EMD-10050), low-pass filtered to 12 Å resolution. Note the P1 and P2 domains of CheA are not present in the monolayer sample. (F) Overlay of the D and E. (G) A slice through a zoomed view of (F), highlighting excess density between CheA P4 domains. (H) Overlay of CSU map from *E. coli* ghost cells with model constructed from 4Q monolayer arrays (PDB 6S1K). For clarity, CheW is not shown.

**Supplementary Figure 4.**
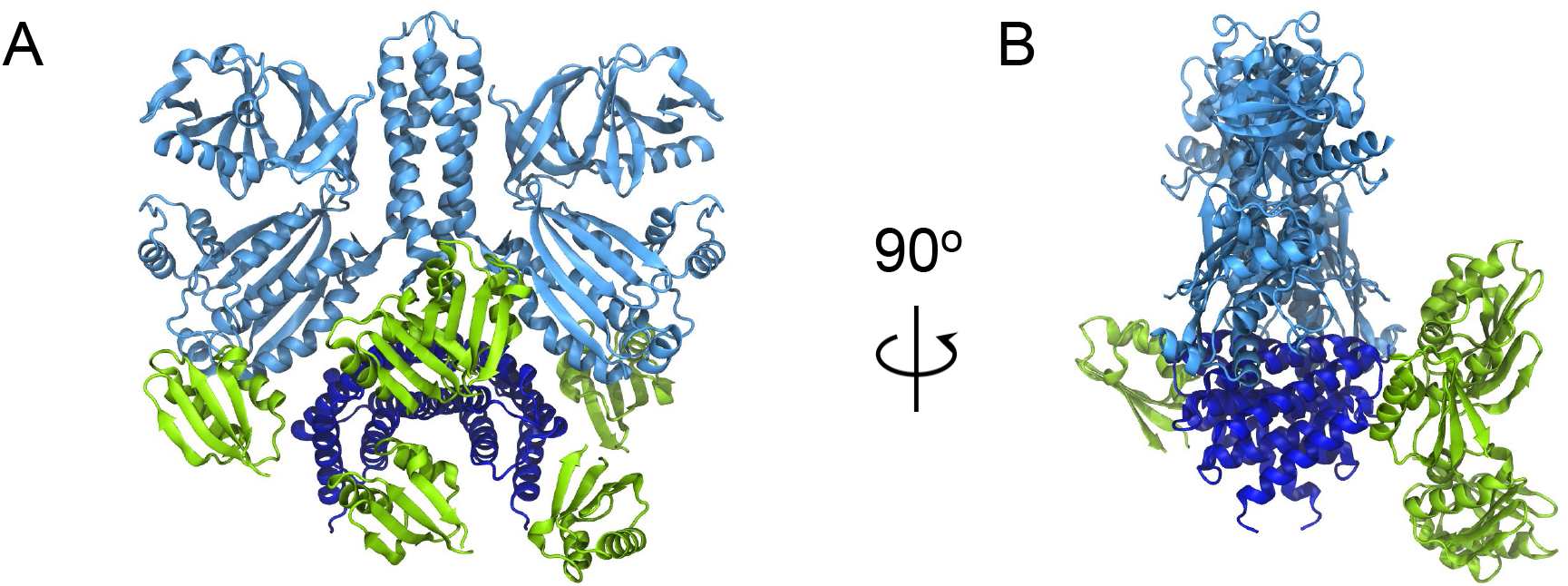
Variable placement of CheA.P2 by AlphaFold2. (A-B) Overlay of five full-length CheA dimer models obtained from Alphafold2, shown from the side (A) and rotated by 90 degrees (B). The position of the P2 domain (green) from one monomer in each model is shown. Note, P2’ (not shown) is placed symmetrically with respect to the opposing side of the CheA dimer. For clarity, CheA.P3-P5 (light blue) and CheA.P1 (dark blue), whose positions are unaltered between the models, are shown here for a single model and the disordered linkers connecting P1 to P2 and P2 to P3 are not shown.

**Supplementary Figure 5.**
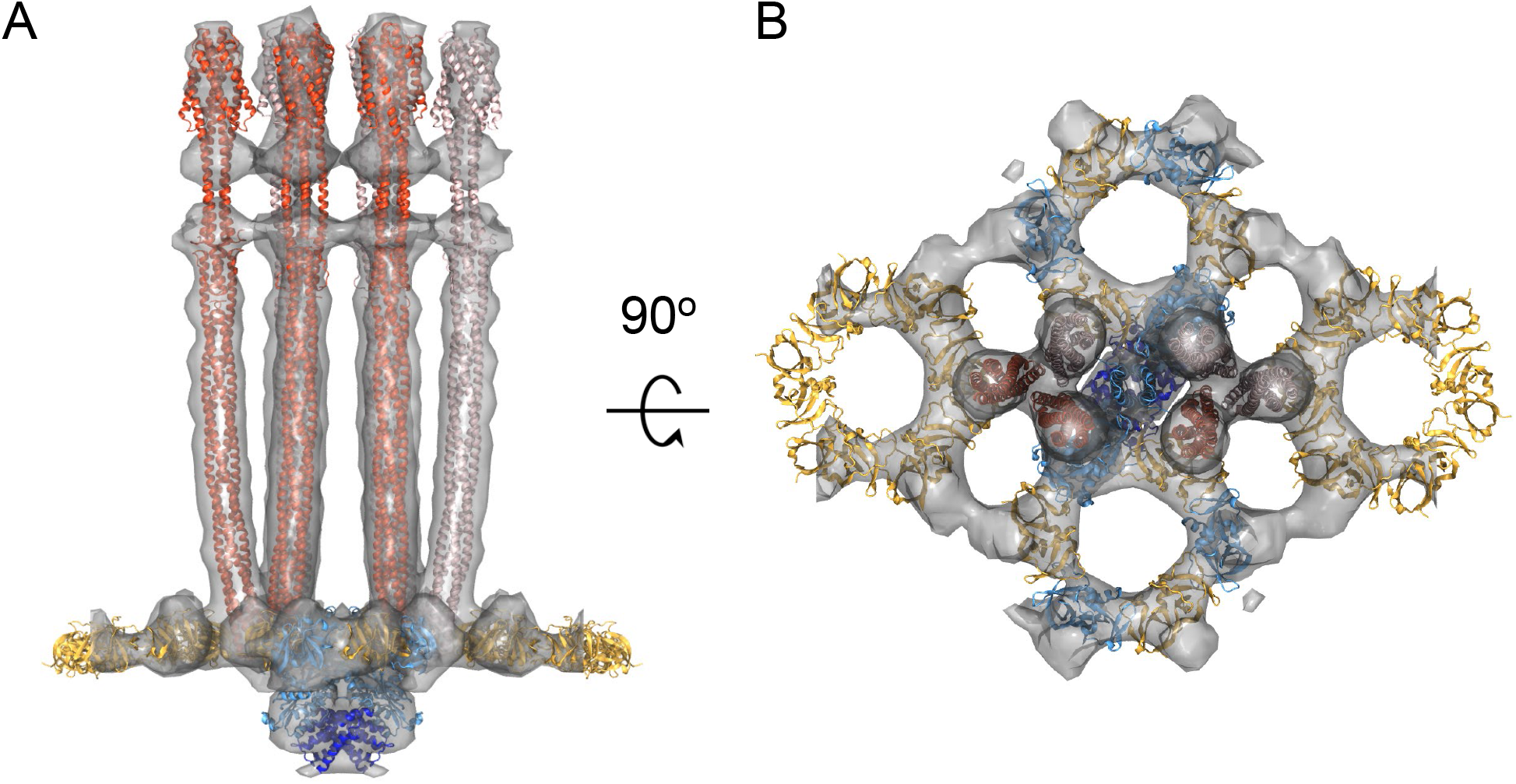
Molecular Dynamics Flexible Fitting (MDFF). (A-B) Overlay between the CSU density map and MDFF-refined CSU model, shown from the side (A) and top (B). The fitted structure included six Tsr homodimers (red/pink), a central CheA.P3-5 dimer (light blue) with associated P1 domains (dark blue), two (CheA.P5/CheW)_3_ rings (blue and gold), and two (CheW)_6_ rings (gold).

**Supplementary Figure 6.**
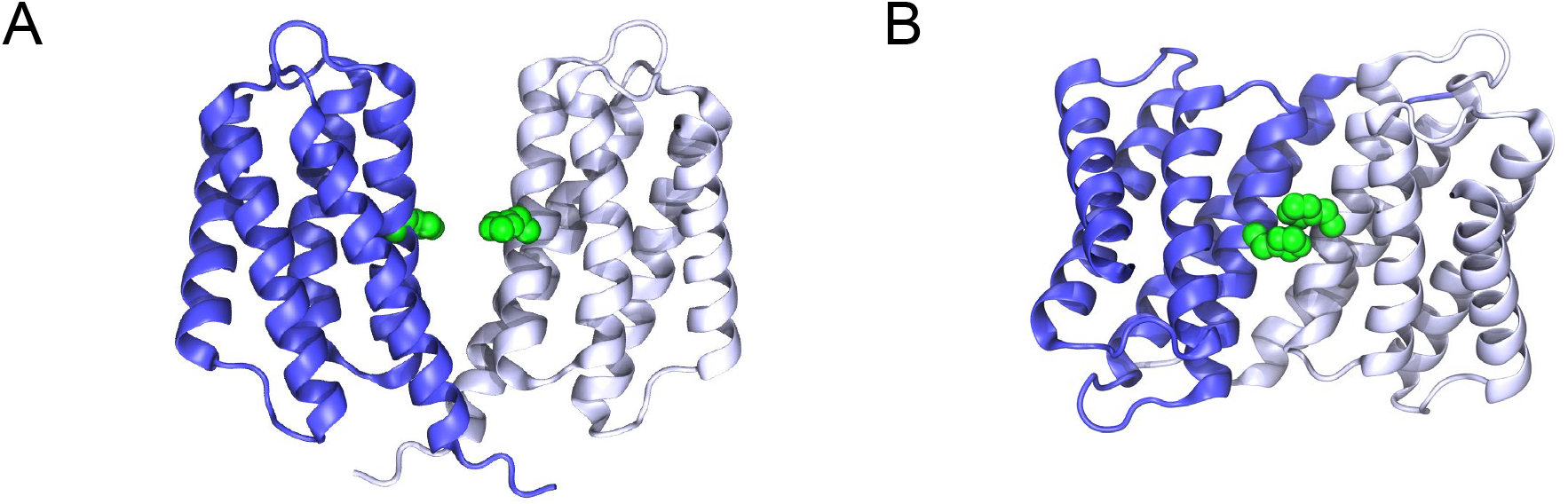
CheA.P1 binding modes predicted by AlphaFold2. (A) Parallel P1 dimer predicted for isolated *E. coli* P1 domains. (B) Anti-parallel P1 dimer predicted for full-length *E. coli* CheA (depicted in Figure 5B). P1 monomers are colored in light and dark blue; the substrate histidine is colored green and shown in VDW representation.

## Movie legends

**Movie 1**. Aligned tilt series of an E-gene lyszed native *E. coli* cell displaying a patch of the chemosensory array lattice.

**Movie 2**. A convolution map resulted from template-matching in emClarity, overlapped with picked subtomograms (green circles).

**Movie 3**. Extract from MD simulation (50-ns) of CSU model in an atomistic lipid bilayer, illustrating interactions between neighbouring receptor ligand binding domains (LBD). Clusters of residues mediating LBD interactions are shown as VDW spheres and coloured in blue (residues E124, K126, R127, D130) or yellow (residues K99, E102, and K103).

